# ExactCN: Predicting Exact Copy Numbers on Whole Exome Sequencing Data

**DOI:** 10.1101/2025.11.24.690086

**Authors:** Erfan FarhangKia, Ahmet Arda Ceylan, Mert Gençtürk, Mehmet Alper Yılmaz, Furkan Karademir, A. Ercument Cicek

**Affiliations:** Department of Computer Engineering, Bilkent University, Ankara, Türkiye

**Keywords:** CNV detection, Deep Learning, Whole Exome Sequencing

## Abstract

The quantification of the precise copy number variations (CNVs) is crucial to understanding the effects of gene dosage, disease severity, and therapeutic response. Although whole-exome sequencing (WES) offers a cost-effective solution for CNV detection in a clinical setting, it introduces several biases, including those related to sequence length, GC content, and the use of targeting probes. Consequently, estimating exact copy numbers remains challenging, especially for WES data. Here, we present ExactCN, a deep learning–based method for estimation of exact copy numbers from WES data per exon. The architecture integrates convolutional layers that extract local read-depth patterns with transformer encoder blocks that capture genomic context and handle sequencing noise. ExactCN is trained on WES samples from the 1000 Genomes Project, using matching WGS-based calls as semi–ground truth. In benchmarks, ExactCN improves the state-of-the-art integer CNV calling performance by reducing the macro-averaged mean absolute error (MAE) from 0.91 to 0.62 and the macro-averaged root mean squared error (RMSE) from 1.31 to 0.78. It also achieves an overall Pearson correlation of 0.669 and Spearman correlation of 0.550, improving the second-best method by 0.641 and 0.482, respectively. Furthermore, a fine-tuned and specialized version of ExactCN for aggregate CNV detection in clinically important duplicated genes *SMN1/2* achieved a macro averaged F1-score of 0.657, and mean absolute error of 0.3. These results substantially improves the state-of-the-art performance and demonstrates the model’s applicability to both research and clinical genomic analyses. ExactCN is available at https://github.com/ciceklab/ExactCN.

## 1 Introduction

Copy number (CN) refers to the number of copies of a particular genomic region present in the DNA of an organism. CNs are typically inferred from sequencing data by analyzing read depth, paired-end mapping discrepancies, or split-read signals [1]. The resulting CN values represent the absolute number of copies or relative abundance compared to a reference. Detection of copy number variants (CNVs), which are structural variations defined by large duplications or deletions in the genome, is critical to understanding genomic variation between individuals and has been implicated in a wide range of genetic disorders, cancers, neuro-developmental conditions, and complex traits such as height, BMI, and drug response [2–5].

Many high-performing computational tools have been developed for CNV detection from whole-genome sequencing (WGS) data [6–22]. Although WGS provides comprehensive genomic coverage, it remains relatively expensive and resource-intensive, limiting its large-scale clinical applications [23, 24]. In comparison, whole-exome sequencing (WES) provides a more cost-effective and interpretable alternative, focusing on protein-coding regions. This makes it a clear solution in the clinic as opposed to WGS. Despite these advantages, CNV detection from WES data has received less methodological attention [8, 25–28, 11, 29] due to biases related to GC content, target length, and mapping efficiency [30, 26]. In this limited scope, categorical detection of CNVs remains the most widely recognized use case. However, accurate estimation of exact copy numbers has broader applications, including gene dosage analysis, evaluation of phenotypic severity, and informed decision-making for therapeutic purposes. For example, in spinal muscular atrophy (SMA), a homozygous deletion of *SMN1* (0 functional copies) causes the severe Type 1 form, requiring immediate therapy, whereas a single-copy state indicates an asymptomatic carrier [31]. Conventional binary “deletion” calls cannot distinguish these states, risking missed interventions and false diagnoses. Similarly, in HER2-positive breast cancer, a high level of *ERBB2* amplification (≥ 6 copies) is associated with significantly improved overall outcomes in trastuzumab-treated patients. In contrast, *PTPN2* gain (CN ≥ 3) is linked to therapeutic resistance and poorer prognosis [32]. Thus, exact copy number genotyping derived from WES read depth can stratify patients for targeted therapy. In pharmacogenomics, *CYP2D6* copy number variation directly influences codeine metabolism and clinical response. Individuals with no functional *CYP2D6* copies (activity score = 0) are poor metabolizers who cannot efficiently convert codeine to morphine, resulting in insufficient analgesia and potential therapeutic failure. In contrast, those carrying more than two functional copies of *CYP2D6* (ultrarapid metabolizers) exhibit excessive morphine formation and severe adverse effects. Therefore, precise determination of *CYP2D6* gene copy number is critical for optimizing codeine dosing and avoiding both undertreatment and toxicity [33]. Finally, in familial risk assessment, distinguishing between 0 and 1 copies in parents enables the accurate calculation of recurrence risk (e.g., 25% for recessive disorders), which guides cascade testing and prenatal counseling. Therefore, methods that output exact copy numbers are not only advantageous but essential for transforming ambiguous CNV calls into actionable clinical insights. There are a few WES-based CNV callers that can make integer CNV calls: CNVKit, Control-FREEC, and GATK-gCNV [27, 11, 8]. However, none of these approaches are learning-based, and we hypothesize that a deep learning-based approach can learn the complex patterns in the noisy read depth signal that indicate a copy number change, given the success of this approach in categorical CNV detection in WES data [25, 26, 15].

Here, we present ExactCN, the first machine learning-based method that accurately infers integer copy numbers at the exon level from WES data. The system model is shown in Figure 1. ExactCN uses convolutional layers and transformer encoders. The model takes as input a read-depth signal with four channels corresponding to bases A, T, C, and G. Then, it integrates convolutional layers for local feature extraction with transformer encoder blocks for contextual learning, followed by a regression head that predicts exact copy numbers at the exome level. We trained ExactCN on 530 WES samples from the 1000 Genomes Project, using high-confidence WGS-derived copy number annotations generated by the DRAGEN pipeline as semi-ground truth.

**Fig. 1:**
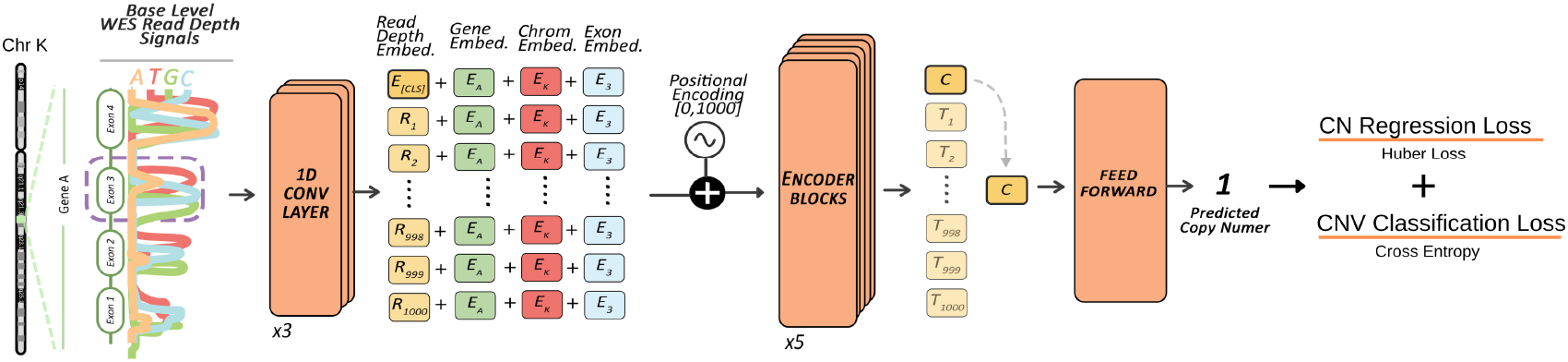
The system model of ExactCN. Base-level whole-exome sequencing (WES) read-depth signals (4 × 1000 tensor for bases A, T, G, C) are processed by three 1D convolutional layers (kernel sizes 9, 9, 1) expanding channels to 64, 128, and 512 features. Each layer is followed by GELU activation, group normalization, and layer normalization. The resulting token sequence is concatenated with a trainable classification token, summed with learned gene, chromosome, and exon embeddings, and then added to sinusoidal positional encoding. This input is fed into a stack of Performer encoder blocks, and the final classification token representation is passed to a two-layer feed-forward MLP with GELU activation to regress the predicted copy number. Training optimizes a composite loss consisting of a Huber regression term and a cross-entropy term to encourage accurate copy-number estimation and correct CNV label classification.

The benchmark results demonstrate that ExactCN achieves superior CNV call performance in integers compared to existing WES-based methods. ExactCN achieves a macro-averaged MAE of 0.622 and RMSE of 0.783, representing improvements of 32% and 40%, over the second-best method, respectively. Examining class-specific performance in regions affected by CNVs, ExactCN demonstrates a 23% improvement in MAE for deletions (0.914 vs 1.181) and a 33% improvement in MAE for duplications (0.948 vs 1.417) compared to the second-best tool for each respective category. ExactCN also demonstrates a substantially higher correlation with ground truth, achieving a macro-averaged Pearson correlation of 0.713, which is substantially higher than the second-best method. To further test the capabilities of ExactCN, we discretize its predictions and benchmark it in the orthogonal task of categorical CNV calling. ExactCN exceeds all competing methods other than the ECOLE, which is the only machine learning-based method. ExactCN achieves a macro-averaged F1-score of 0.722, trailing ECOLE-FT by only 0.009 (0.731 vs 0.722). This represents a 49% improvement over the next best method, which achieves 0.485. Notably, ExactCN also achieves the highest macro-averaged precision (0.822) and demonstrates superior class-specific precision for both deletions (0.755) and duplications (0.713).

Finally, we introduce ExactCN^*SMN*^, a *SMN* -specific CNV caller that is a fine-tuned derivative of ExactCN, achieving a macro-averaged F1-Score of 0.657 (an improvement of 0.253 over the second-best method), with balanced precision and recall. To our knowledge, it is the first WES-based CNV caller that can make integer CNV calls for this gene, representing a proof-of-concept effort to allow dose-specific and personalized treatment for SMN patients. The code is available at https://github.com/ciceklab/ExactCN.

## 2 Materials and Methods

### 2.1 Problem formulation

ExactCN estimates exact copy numbers for each exon. The observed data comprise a read-depth tensor 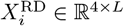 with four channels corresponding to bases A, T, C and G, padded to a fixed length *L* = 1000. Entry 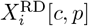 counts the number of sequencing reads supporting base *c* at position *p*. To remove global coverage variation, we subtract a channel-specific mean *µ*_*c*_ and divide by a channel-specific standard deviation *σ*_*c*_, calculated from the training cohort. Let *ℓ*_*i*_ ≤ *L* denote the true exon length in base pairs. We place the *ℓ*_*i*_ observed bases at the *right* end of the fixed-length tensor and pad the remaining *L* − *ℓ*_*i*_ leading positions with a constant value as shown in Eq.(1).

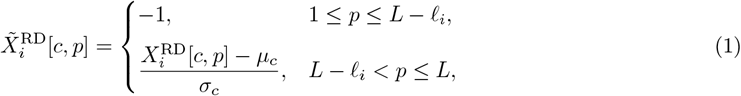

Thus, the leading padded positions encode non-informative context with the constant value −1, while the remaining positions store the normalized exon signal.

We use the integer copy numbers *y*_*i*_ ∈ ℤ_*≥*0_ obtained from the matched high-coverage WGS via Illumina DRAGEN CNV pipeline as the semi-ground truth regression targets. To obtain the CNV labels, we convert these integer copy numbers into a tri-class label *t*_*i*_ *∈* {0, 1, 2} by grouping *y*_*i*_ *∈* {0, 1} as deletion (*t*_*i*_ = 0), *y*_*i*_ = 2 as no-call (diploid state) (*t*_*i*_ = 1), and *y*_*i*_ *≥* 3 as duplication (*t*_*i*_ = 2). For training, we utilize the equivalent one-hot encoded vector **t**_*i*_ *∈* {0, 1}^3^. Each exon *i* is defined by its chromosome index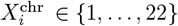, genomic coordinates 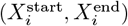, gene identifier *g*_*i*_, and exon index *e*_*i*_. We learn a mapping *f*_*θ*_ that takes as input the normalized tensor 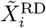 together with its metadata 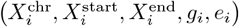 and outputs the continuous prediction *ŷ*_*i*_. The parameters *θ* are fitted by minimising the composite loss over N training exons:

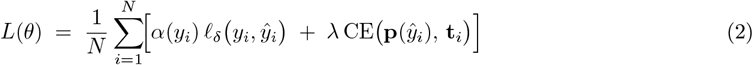

where *ℓ*_*δ*_ is the Huber loss term, *α*(*y*_*i*_) assigns weights (4, 8, 4) to deletions, no-call exons, and duplications, respectively. CE is the cross-entropy term, using class weights of 1 for deletions, 1 for no-call exons, and 10 for duplications, and *λ* = 0.9 balances the regression and classification objectives. This combined loss is natural because ExactCN simultaneously performs regression, estimating a continuous copy number, and classification, assigning each exon to the deletion, no-call, or duplication class. The Huber term encourages *ŷ*_*i*_ to match the discrete DRAGEN count *y*_*i*_, while the class-weighted cross-entropy uses **p**(*ŷ*_*i*_) (see Eq. (3)) to ensure that the predicted value falls decisively into one of the three states. To compute the cross-entropy term CE, we first evaluate three sigmoid-based scoring functions:

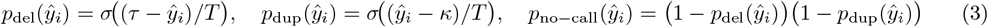

where *σ*(*z*) = 1*/*(1 + *e*^−*z*^) is the logistic function, *τ* = 1.6 and *κ* = 2.2 are the deletion and duplication thresholds, and *T* = 0.1 is a temperature parameter. We then normalise these scores to sum to one to obtain the softly binned probability vector **p**(*ŷ*_*i*_) = *p*_del_, *p*_no−call_, *p*_dup_.

### 2.2 Model description

The architecture of ExactCN is shown in Fig. 1. For each exon i, the model takes a normalized read-depth tensor 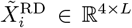 of fixed length 1000 with four channels corresponding to bases A, T, C, and G as input. Three consecutive one-dimensional convolution layers with kernel sizes 9, 9, and 1 expand the four input channels to 64, 128, and finally *D* = 512 feature dimensions. Each convolution is followed by a GELU activation, group normalization across channels, and layer normalization across tokens.

Following the convolutional layers, a trainable classification token *z*_cls_ ∈ ℝ^*D*^ is prepended to the sequence of token embeddings. To incorporate genomic context, we introduce three learnable embeddings representing the gene identifier, chromosome index, and exon index. Their sum yields a context vector *c*_*i*_ ∈ ℝ^*D*^, which is added element-wise to each token in the sequence. Finally, to encode absolute position, we add a fixed sinusoidal positional embedding *E*^pos^ ∈ ℝ ^(*L*+1)*×D*^ to the resulting sequence. For a position *p*∈ {0, …, *L*} and even/odd feature indices 2*j* and 2*j* + 1, we define:

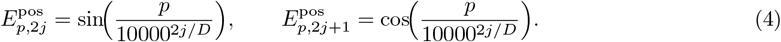

The final input representation is processed by a stack of *M* = 5 Performer encoder layers, an efficient variant of the Transformer that uses kernel approximation to scale linearly in sequence length [34]. Each layer applies layer normalization, multi-head Performer attention, residual connections, and a two-layer feed-forward network with GELU activation. We mask padded positions in the attention computation so they do not influence the context. Let 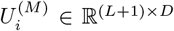 be the output of the final Performer layer. The first row *u*_*i*,cls_ ∈ ℝ^*D*^ corresponds to the classification token, which aggregates information across the exon. A two-layer multilayer perceptron then maps this vector to the predicted copy number:

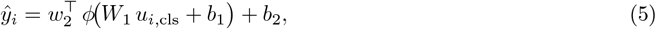

where *W*_1_ ∈ ℝ^*D×D*^ and *w*_2_ ∈ ℝ^*D*^ are weight matrices, *b*_1_ ∈ ℝ^*D*^ and *b*_2_ ∈ ℝ are biases, and *ϕ* is the GELU activation. The model is trained end-to-end using the composite loss in Eq. (2).

### 2.3 Comparison with the existing methods

ExactCN is the first machine learning–based approach tailored to estimation of integer copy numbers in exonic DNA regions. On the other hand, there are prior methods addressing similar problems, but they were developed with different focuses and underlying principles. For instance, CNVnator segments read-depth histograms computed over fixed bins using a mean-shift algorithm augmented with multi-bandwidth partitioning and GC-content correction [7]. Although it is sensitive, its resolution is constrained by the selected bin size. Moreover, since it is specialized for WGS, the method performs poorly when applied to WES due to inherent biases in the data. GATK-gCNV combines negative-binomial factor analysis of read counts with a hierarchical hidden Markov model to jointly detect and genotype CNVs; this Bayesian model can recover rare events in large cohorts but requires substantial computational resources [11]. CNVkit integrates both on-target and off-target reads from hybrid capture, corrects biases due to GC content and repetitive sequences, and segments normalized log2 copy ratios using circular binary segmentation, yielding categorical copy number calls [27]. Control-FREEC constructs raw copy number profiles using non-overlapping read-count windows, normalizes them via GC content or a matched control sample, applies LASSO-based segmentation, and assigns integer copy number states; it can operate without controls but was designed primarily for WGS data [8]. Finally, ECOLE employs a transformer encoder to classify exons into deletion, neutral, or duplication categories based on WES read-depth signals; it does not regress continuous copy numbers [25]. By regressing integer copy numbers with a linear-time transformer, ExactCN provides a unified framework for both CNV detection and precise CN estimation.

### 2.4 Datasets

#### Autosomal Exome Dataset

We assembled a cohort of WES samples with matching high-coverage WGS data from the 1000 Genomes Dataset [35]. We randomly selected a total of 687 WES samples drawn from 26 populations and partitioned them into 530 training samples, 147 test samples, and 10 validation samples. The sample IDs we used are listed in Supplemental Tables 11, 12, and 13. For each sample, we computed four-channel read-depth vectors across 245,965 autosomal exons, normalized them, and paired them with integer copy number labels derived from the matched WGS reads using the Illumina DRAGEN CNV pipeline as semi-ground truth.

Across the 530 training samples, we analysed 98,197,340 autosomal exons. Of these, 155,306 exons were labeled as deletions and 115,850 as duplications, highlighting the extreme class imbalance in the training set. The 147-sample test set contained 27,235,866 exons, comprising 44,352 deletions (CN ≤ 1), 40,298 duplications (CN ≥ 3), and 27,151,216 diploid regions. The 10-sample validation set comprised 1,852,779 exons with 2,691 deletions, 1,567 duplications, and 1,848,521 diploid exons. These counts correspond to deletion frequencies of approximately 0.16% and duplication frequencies of 0.08–0.15% in the held-out subsets, highlighting the extreme class imbalance characteristic of copy number studies.

#### *SMN* Dataset

The *SMN* dataset is used to evaluate ExactCN on a targeted CNV prediction task for the *SMN1* and *SMN2* genes. The high sequence similarity between *SMN1* and *SMN2* causes WES reads to align ambiguously. As a result, default DRAGEN calls often return no-call for these loci. To obtain reliable semi-ground-truth labels for *SMN1/2*, we used the SMNCopyNumberCaller [36]. It is a tool specialized for estimating the copy numbers of *SMN1* and *SMN2* using WGS data, which is substantially more reliable than WES. Additionally, since the depth signal cannot be assigned to either gene with confidence, we focused on aggregate copy-number estimation by summing read-depth across both loci to produce a single combined signal.

The dataset comprises 95 whole-exome sequencing samples from the 1000 Genomes Project, with a focus on the *SMN1* and *SMN2* loci. To maintain consistency with the pre-training setup, we preserved the same train–test split for the deletion and no-call labels: the 55 samples used for training during pre-training remained in the training set, and the corresponding 40 samples were retained as the test set. During dataset curation, two duplication samples were excluded as outliers based on a mean read-depth threshold of 10. The final *SMN* test set contains 40 samples (9 duplications, 17 no-call, 14 deletions), while the fine-tuning set comprised 55 samples (14 duplications, 20 deletions, 21 no-call). The list of sample IDs are provided in Supplementary Table 15.

## 3 Results

### 3.1 Experimental Setup

#### Compared Method

We compare ExactCN with five WES-based CNV callers: CNVkit, GATK-gCNV, Control-FREEC, and ECOLE. To ensure a fair comparison with ExactCN, we also fine-tuned ECOLE on the same high-quality DRAGEN-labeled dataset as ExactCN, referred to as ECOLE-FT.

CNVkit, GATK-gCNV, and Control-FREEC output integer copy numbers, which we evaluate for both regression and classification performance. ECOLE outputs categorical CNV predictions (deletion, no-call, duplication); therefore, both ECOLE and ECOLE-FT are evaluated only for classification performance. ECOLE was originally trained on CNVNator-derived semi–ground truth labels, which differ from the DRAGEN-based semi-ground truth labels used for training ExactCN. To ensure a fair comparison, we fine-tuned ECOLE on samples annotated with DRAGEN semi–ground–truth labels, and this version is referred to as ECOLE-FT. Fine-tuning was performed on 65 training samples for one epoch on a NVIDIA TITAN RTX GPU using default parameters (See Supplementary Table 14). The sample IDs included in the fine-tuning set are listed in Supplementary Table 14. The samples, ground truth data, and CNV predictions to reproduce the analyses can be accessed through https://zenodo.org/records/17666701.

#### Parameter Settings

We run GATK-gCNV cohort analysis using 147 test samples simultaneously, with default parameters and internal GC content normalization. For CNVKit, we generated a pooled reference from all samples and used that as a reference sample in the analysis, with default hyperparameters. We ran the Control-FREEC with GC normalization and recommended parameters (See Supplementary Table 10). ECOLE and ECOLE-FT were also run using their default parameter settings (See Supplementary Table 9).

#### Training ExactCN

ExactCN is trained on 530 whole-exome sequencing samples drawn from the 1000 Genomes Project. For each sample, we extract four-channel read-depth profiles at 245,965 autosomal exons and pair them with integer copy number labels inferred from the matched high-coverage whole-genome sequencing data using the DRAGEN CNV pipeline. The resulting training set exhibits extreme class imbalance, with fewer than one in a thousand exons harboring a deletion or duplication. To prevent the model from being overwhelmed by the diploid background, mini-batches of 256 exons are drawn without replacement using a stratified sampler that guarantees at least one deletion and one duplication every eight batches, while preserving the overall class proportions. Each exon is normalized by channel-specific means and standard deviations computed from the training cohort, and masked positions beyond the true exon length are set to a constant.

During optimisation, we minimise a composite objective that combines a weighted Huber loss with a soft-binned cross-entropy loss. The Huber term uses parameter *δ* = 2 and class weights (8, 4, 4) for the no-call, deletion, and duplication categories, respectively, reflecting their relative frequencies. The cross-entropy term employs class weights (1, 1, 10), discretises the regressed copy numbers into deletion, neutral, and duplication probabilities using thresholds *τ* = 1.6 and *κ* = 2.2, and applies a temperature *t* = 0.1 to control softness; a scaling factor *γ* = 0.9 balances this term against the Huber loss.

Implementation is carried out in PyTorch. The network integrates three one-dimensional convolutions, five linear-time Performer encoder layers with eight attention heads and a hidden size of 512, as well as a two-layer regression head. Model parameters are optimised with the AdamW algorithm using an initial learning rate of 1 *×* 10^−4^, weight decay of 0.01, and a cosine annealing learning rate scheduler. Training is performed for ten epochs on a single NVIDIA TITAN RTX GPU with 24 GB of memory; under this configuration, the model converges in roughly 168 hours. After each epoch, we evaluate the network on a validation set of 10 WES samples, adjust the class weights and decision thresholds based on the validation performance, and retain the checkpoint that yields the lowest mean absolute error.

#### Training ExactCN^*SMN*^

ExactCN^*SMN*^ represents a fine-tuned derivative of ExactCN, optimized to regress exact aggregate integer copy numbers for *SMN1/2* loci. The model’s weights are initialized from the pretrained ExactCN model parameters and fine-tuned for 1500 epochs on an NVIDIA TITAN RTX GPU. ExactCN^*SMN*^ is trained on 50 WES samples that consist of 13 duplications, 16 deletions, and 21 no-call regions. The general fine-tuning procedure was identical to that used for training ExactCN, including the pre-processing of read-depth profiles at the exon level, model architecture, loss formulation, and the learning rate schedule. Since the fine-tuning dataset was smaller and its class composition differed from the original training cohort, no stratified sampler was used. Instead, the class weights in the cross-entropy loss were redefined to match the distribution of the fine-tuning set, with weights of 1 for deletions, 4 for no-call regions, and 2 for duplications. All remaining hyperparameters were kept unchanged from the original setup.

### 3.2 Evaluation criteria

We evaluate ExactCN ‘s performance across two tasks: exact copy-number regression and CNV detection. For the regression task, we report the Mean Absolute Error (MAE), Root Mean Squared Error (RMSE), and the Pearson and Spearman correlations, both overall and segmented by copy-number categories representing CNVs. The threshold values for this categorization were determined based on performance analysis using the validation set: deletions (CN ≤ 1.7), no event (1.7 *<* CN *<* 2.3), and duplications (CN ≥ 2.3) (see Supp. Fig. 1). This categorization is also used for the CNV detection task, for which we report precision, recall, and F1-scores separately for deletions and duplications. CNVkit, GATK-gCNV, and Control-FREEC output integer copy numbers; we evaluate them on both tasks, by categorizing integer copy number 2 as no-call, 0 and 1 as deletion, and more than 3 as duplication. ECOLE and ECOLE-FT are evaluated only on CNV detection.

For the evaluation of ExactCN^*SMN*^, we apply the same categorization approach and thresholds. ExactCN^*SMN*^ is assessed solely on the CNV detection task, and only within the *SMN1/2* region in an aggregated manner.

### 3.3 The Performance of ExactCN on Integer Copy Number Prediction

We evaluated ExactCN on a held-out test set of 147 WES samples with semi-ground truth copy number labels derived from matching high-coverage WGS data using the Illumina DRAGEN CNV pipeline. For regression evaluation, we rounded ExactCN’s continuous copy number predictions to the nearest integer, since all compared methods output integer predictions. ExactCN shows superior regression performance across all evaluated metrics (Figure 2). For overall performance, ExactCN achieves an MAE of 0.003 and an RMSE of 0.079, representing a 14-fold improvement in MAE over the next-best method, CNVkit (MAE = 0.049). Given the severe class imbalance in genomic data (deletion: 0.16%, duplication: 0.15%, no-call: 99.69%), macro-averaged metrics provide an unbiased evaluation across all region types. ExactCN achieves macro-averaged MAE of 0.620 and RMSE of 0.781, representing 32% and 40% improvements over the next best method.

**Fig. 2:**
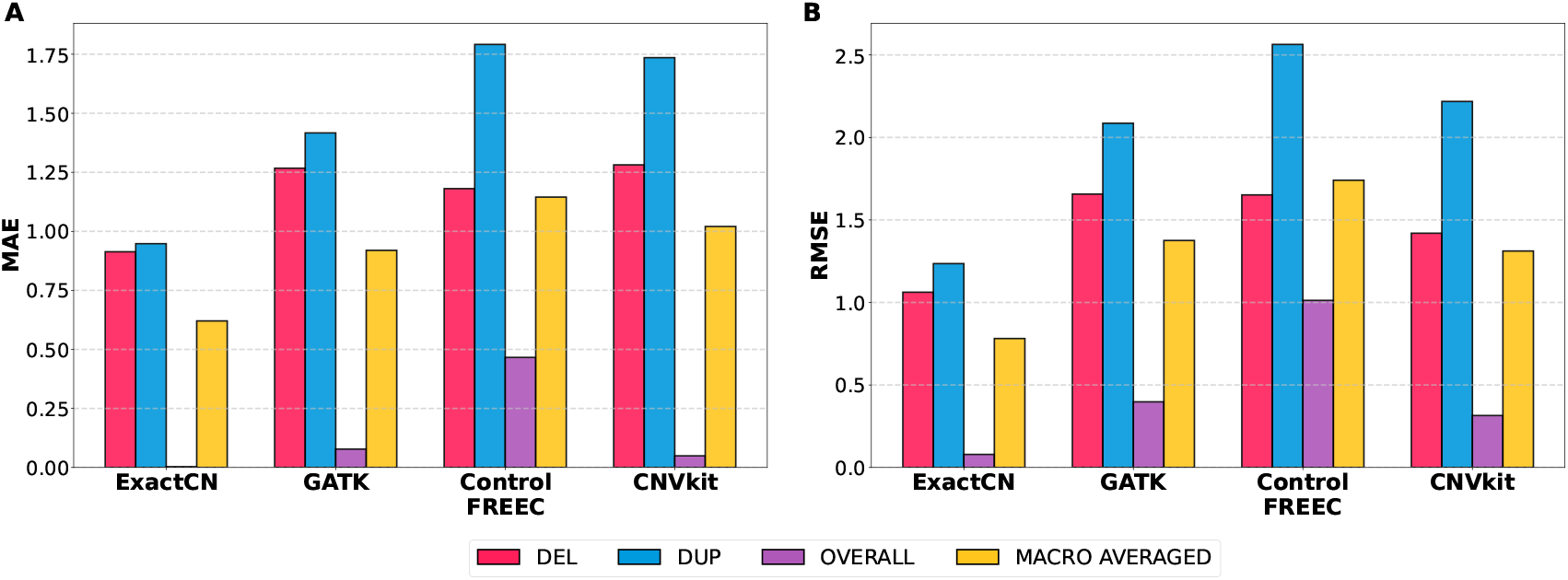
Comparative evaluation of CNV caller performance using mean absolute error (MAE) and root mean square error (RMSE). Four tools were assessed on deletion (DEL), duplication (DUP), and no-call regions using DRAGEN-derived semi-ground-truth labels from matched WGS samples. Overall metrics reflect performance across all regions, while macro-averaged metrics treat all classes equally, providing unbiased evaluation under severe class imbalance (DEL: 0.16%, DUP: 0.15%, no-call: 99.69%). **A**. MAE results for each variant type, overall error, and macro-averaged error across tools. **B**. RMSE results for each variant type, overall error, and macro-averaged error across tools.

Examining class-specific performance in CNV-affected regions, ExactCN maintains consistently low error rates across all classes. For deletion regions, ExactCN achieves an MAE of 0.914 and an RMSE of 1.062, representing 23-25% improvement over the next-best method. For duplication regions, ExactCN shows even more substantial gains with an MAE of 0.953 and an RMSE of 1.239, achieving 32-40% improvements over the next-best method. The relatively higher error in duplications compared to deletions reflects the inherent challenge of distinguishing between CN = 3, CN = 4, and higher-order amplifications from noisy WES data. For no-call regions (CN = 2), ExactCN achieves exceptionally low error (MAE = 0.0005, RMSE = 0.046), demonstrating its ability to accurately distinguish diploid states from true CNVs. Finally, we show the confusion matrices in Figure 3. We observe that ExactCN shows the strongest diagonal pattern, indicating a higher correlation with the ground truth and demonstrating a higher prediction accuracy. GATK does not make CN predictions above 5, which leads to underestimating many CNVs, whereas ExactCN has learned that CNs over 10 are very rare and minimizes that error. Control-FREEC is very liberal and makes calls in the full range, but this leads to many overestimations. Additional regression performance analyses are presented in Supplementary Tables 3 and 4.

**Fig. 3:**
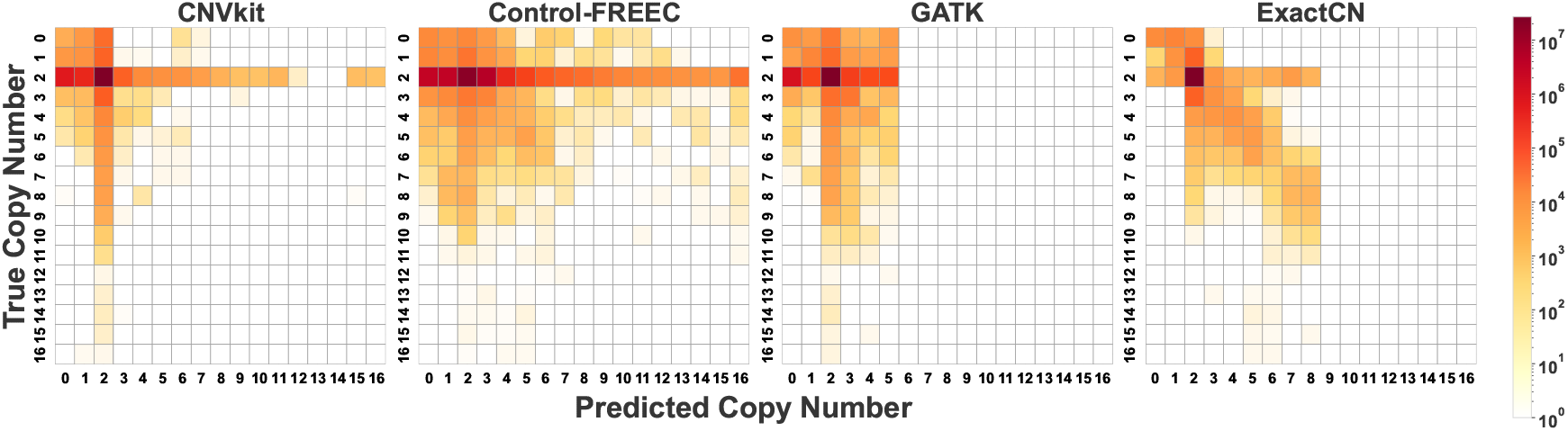
The confusion matrices comparing copy number predictions of CNVKit, Control-FREEC, GATK, and ExactCN. Rows represent true copy numbers, columns represent predicted copy numbers (0-16, with predictions >16 capped at 16). True copy numbers gathered using DRAGEN from matched WGS samples. Color intensity (log scale) indicates prediction count. ExactCN shows the strongest diagonal pattern, demonstrating higher prediction accuracy.

### 3.4 The Performance of ExactCN on Categorical CNV Prediction

Although ExactCN was designed for exact copy number regression rather than categorical CNV detection, its continuous predictions can be effectively converted to CNV calls using the thresholds defined above. Figure 4 shows the classification performance across all methods.

**Fig. 4:**
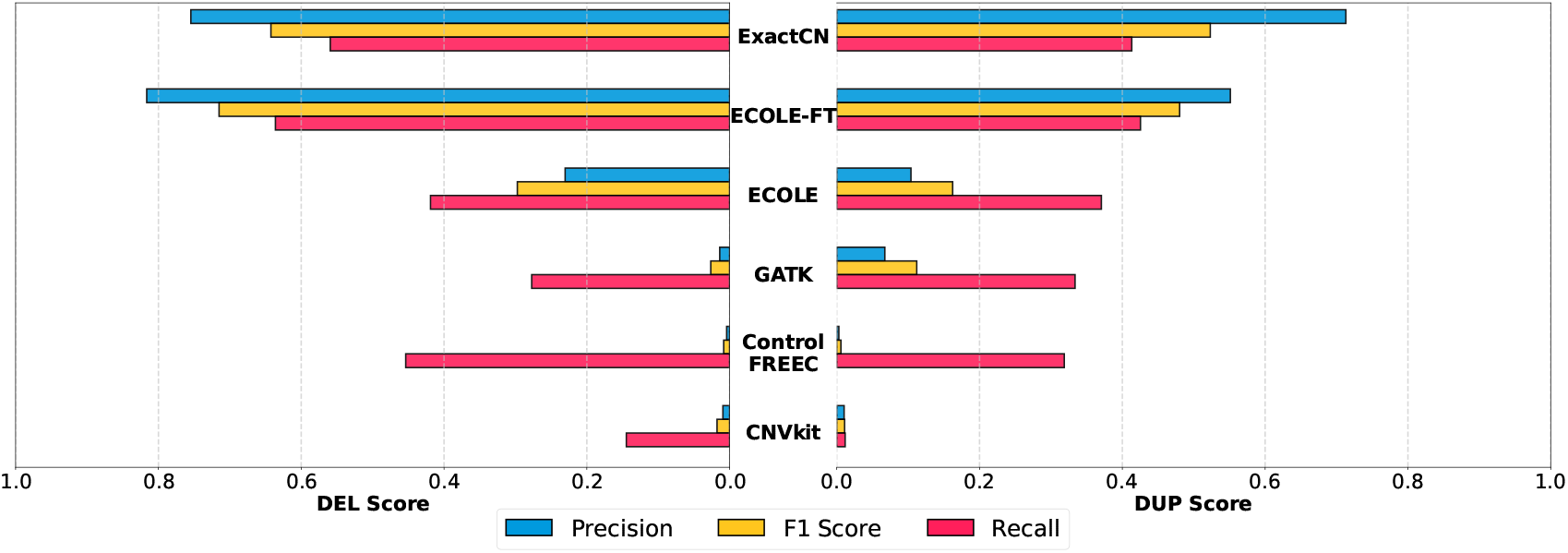
Left: Precision, recall, and F1-scores for deletion detection across tools. Right: Precision, recall, and F1-scores for duplication detection across tools. ECOLE-FT is fine-tuned on the same high-quality DRAGEN reference set used to train ExactCN. CNVkit and Control-FREEC output integer copy number predictions, which are discretized into deletion, duplication, and no-call. ExactCN, originally designed for continuous copy number estimation, is also discretized into the same categories.

For deletion detection, ExactCN achieves an F1-score of 0.642, striking a balance between high precision (0.755) and moderate recall (0.559). Notably, ExactCN achieves the second-highest precision among all methods for deletion detection, surpassed only by the fine-tuned ECOLE model. Compared to the pre-trained ECOLE model, trained on the 1000 Genomes dataset with semi-ground-truth CNVNator labels, ExactCN demonstrates a 3.3-fold improvement in precision and a 2.2-fold improvement in F1-score. To ensure a fair comparison, we fine-tuned ECOLE on the same high-quality DRAGEN reference set used to train ExactCN, named ECOLE-FT. This fine-tuned version achieves the highest F1-score (0.715) with improved recall (0.636) and superior precision (0.816) compared to ExactCN. While ECOLE-FT demonstrates 8% higher precision, ExactCN maintains competitive performance with substantially higher precision than the pretrained ECOLE baseline, demonstrating the effectiveness of our approach for achieving high specificity in CNV detection without task-specific fine-tuning.

For duplication detection, ExactCN achieves an F1-score of 0.523 with a precision of 0.713 and a recall of 0.413, demonstrating 6.9-fold and 3.2-fold improvements in precision and F1-score, respectively, over the pretrained ECOLE, and 29% and 9% improvements over ECOLE-FT. At the macro-averaged level, ExactCN achieves the highest precision (0.822), representing a 4% improvement over ECOLE-FT and 1.9-fold improvement over pretrained ECOLE, while ECOLE-FT achieves a 1.4% higher macro-averaged F1-score (0.731 vs. 0.722) due to its higher recall (0.687 vs. 0.657 for ExactCN). Extended categorical CNV prediction performance analyzes and corresponding confusion matrices are presented in Supplementary Tables 1, 2 and 7.

Traditional CNV callers show lower performance in both categories. GATK-gCNV achieves F1-scores of 0.026 for deletions and 0.112 for duplications, while having moderate recall values. Control-FREEC and CNVkit show F1-scores of less than 0.02 for both classes. Although Control-FREEC achieves a relatively high recall (0.454 for deletions), its low precision (0.004) results in more false positive calls, making it less suitable in practice. These results demonstrate that ExactCN successfully addresses both use cases: it provides state-of-the-art copy number estimates while maintaining competitive performance for categorical CNV detection, with particular strength in precision.

### 3.5 Copy Number Variation Prediction Performance of ExactCN^*SMN*^ on *SMN1/2*

The *SMN1/2* genes are clinically critical duplicated genomic regions whose extreme sequence similarity poses substantial challenges for copy-number analysis. These challenges are amplified in standard whole-exome sequencing (WES), where most exons lack reliable mappability, and the noisy read depth and coverage variability make accurate CNV prediction particularly difficult.

Because of these challenges, conventional WES-based CNV callers that are not specifically designed for *SMN1/2* often produce highly imbalanced predictions at this locus. As shown in Table 2, tools such as ExactCN, CNVkit, Control-FREEC, ECOLE, and ECOLE-FT display a strong bias towards the no-call state. Although no-call recall values range from 0.7 to 1.0, they yield negligible or even zero recall for both deletion and duplication events. With the exception of GATK, most tools effectively default to predicting no-call for the vast majority of samples (See Supplementary Table 8 for the corresponding confusion matrices).

**Table 1:** Correlation and agreement metrics for exact copy number estimation on 147 WES samples. Pearson correlation measures linear relationships, Spearman correlation measures monotonic relationships, and Quadratic Weighted Kappa (QWK) measures ordinal agreement. Bold values indicate the best performance in each column. DRAGEN-based calls are used as the semi-ground truth to calculate the metrics.

| Tools | Pearson Correlation |  |  | Spearman Correlation |  |  | QWK |
| --- | --- | --- | --- | --- | --- | --- | --- |
|  | DEL | DUP | Overall | DEL | DUP | Overall |  |
| CNVkit | -0.087 | 0.049 | 0.014 | -0.069 | 0.030 | 0.024 | 0.014 |
| Control-FREEC | 0.063 | 0.204 | 0.028 | 0.094 | 0.184 | 0.021 | 0.005 |
| GATK | 0.058 | -0.053 | 0.032 | 0.041 | -0.193 | 0.068 | 0.034 |
| ExactCN | **0.642** | **0.783** | **0.669** | **0.650** | **0.762** | **0.550** | **0.510** |

**Table 2:** Aggregated *SMN1/2* CNV calling performances of tools on 40 WES samples from the 1000 Genome Project. Chen et al. calls are used as the semi-ground truth to calculate the metrics.

| Tool | DEL<br>Precision | DUP<br>Precision | NO-CALL<br>Precision | DEL<br>Recall | DUP<br>Recall | NO-CALL<br>Recall | DEL<br>F<sub>1</sub> | DUP<br>F<sub>1</sub> | NO-CALL<br>F<sub>1</sub> |
| --- | --- | --- | --- | --- | --- | --- | --- | --- | --- |
| GATK | 0.143 | 0.250 | 0.381 | 0.071 | 0.333 | 0.471 | 0.095 | 0.286 | 0.421 |
| Control-FREEC | 0.600 | 0.200 | 0.480 | 0.214 | 0.222 | 0.706 | 0.316 | 0.211 | 0.571 |
| CNVkit | **1.000** | **1.000** | 0.447 | 0.071 | 0.111 | **1.000** | 0.133 | 0.200 | 0.618 |
| ECOLE | **1.000** | 0.000 | 0.472 | 0.286 | 0.000 | **1.000** | 0.444 | 0.000 | 0.642 |
| ECOLE-FT | 0.000 | 0.000 | 0.425 | 0.000 | 0.000 | **1.000** | 0.000 | 0.000 | 0.596 |
| ExactCN | 0.000 | 0.000 | 0.425 | 0.000 | 0.000 | **1.000** | 0.000 | 0.000 | 0.596 |
| ExactCN<sup>SMN</sup> | 0.786 | 0.800 | **0.667** | **0.786** | **0.444** | 0.824 | **0.786** | **0.571** | **0.737** |

Overall, these patterns demonstrate that conventional WES-based CNV callers fail to capture the true dynamics of *SMN1/2* copy-number variation and tend to collapse their outputs into a single dominant call type. This underscores the need for specialized, *SMN* -focused CNV detection methods.

To address this gap, we fine-tuned ExactCN using only CNV calls for *SMN* in 50 WES samples and introduced ExactCN^*SMN*^. For each sample, the read-depth profiles of *SMN1* and *SMN2* were summed to form an aggregate signal.Given the extensive sequence homology between *SMN1* and *SMN2*, reliable coverage in our WES data is restricted to Exon 7. This region is critical because it harbors the specific base difference (*c*.840*C > T*) that disrupts a splicing enhancer, distinguishing the *SMN1* gene from the *SMN2* paralog [36]. To the best of our knowledge, ExactCN^*SMN*^ is the first CNV method tailored specifically for integer CNV detection on *SMN1/2* using WES data.

The evaluation was performed on 40 WES test samples from the 1000 Genomes Project. ExactCN^*SMN*^ demonstrates superior regression performance with an overall MAE of 0.3 (an improvement of 0.119 over the next best method) and a RMSE of 0.592 (an improvement of 0.128 over the next best method), out-performing the baseline tools in the deletion, duplication, and no-call regions (see Supplementary Table 6). Note that the next best method in both categories is ExactCN without fine-tuning. We also compared all methods in categorical CNV detection after discretizing the calls of ExactCN and ExactCN^*SMN*^. In this scenario, F1-scores of our model are 0.786 for deletions, 0.571 for duplications, and 0.657 overall (See Table 2 and Supplementary Table 5). These correspond to 0.268, 0.36, and 0.253 improvements over the next best methods, ECOLE and CNVkit, respectively. Overall, these results show that ExactCN^*SMN*^ achieves accurate CNV detection in the challenging, highly duplicated *SMN* region.

## 4 Discussion

The results of this study reveal the efficacy of our deep learning-based exact copy number estimator, ExactCN, in addressing the challenges associated with WES data analysis, particularly in estimating precise copy numbers and detecting copy number variations from noisy exome sequencing data. The use of such a complex neural network architecture is justified by examples in the literature, which clearly show that learning CNV patterns in high-throughput sequencing data requires deep architectures [15, 12], especially when working with WES data [25]. As demonstrated, ExactCN achieves substantial improvements over existing CNV calling methods, outperforming other CNV callers, including CNVkit, GATK-gCNV, and Control-FREEC, in both exact copy number regression and categorical CNV detection tasks. Furthermore, despite not being explicitly optimized for categorical CNV calling, ExactCN achieves competitive performance with the fine-tuned ECOLE-FT model, while demonstrating superior precision and substantially outperforming the pretrained ECOLE baseline. Additionally, we introduce ExactCN^*SMN*^, a fine-tuned variant of ExactCN that represents the first dedicated *SMN* -specific WES caller reported in the literature.

One concern for a learning-based method such as ExactCN is that the model may memorize the frequency of CNVs in each exon and fix the labels at test time. This would mean that it does not learn the relationship between the read depth signal and the labels, but samples the calls based on the gene index. We observe that this is not the case. First, we observe that only %18 of exons with deletion calls and 32% of exons with duplication calls in the training set also have events in the test set. This indicates that test and training events are largely distinct. We also observe that ExactCN makes calls for exons for which there are no events in the training data, but do so in the test data. Finally, we utilized the fine-tuning process of ExactCN^*SMN*^ to establish a more challenging scenario for model evaluation and to assess the risk of label fixing. We included six exon samples from the pre-training dataset, which were labeled as no-call by DRAGEN, into the *SMN* test set. These samples are labeled with a duplication event by the SMNCopyNumberCaller. The model *knows* these samples, so it should report no-call if memorized; otherwise, if fine-tuning the model on *SMN* -specific signals. Indeed, the model perfectly captures all these events even though it was *misguided* by the pre-training. This result shows that model does not memorize the calls and can be easily configured for target genes of clinical interest with a small number of samples.

Despite these promising results, some limitations remain. ExactCN has only been trained using short-read data, and therefore is not applicable to long-read, single-cell, or similar other technologies. Another limitation of ExactCN is its inability to detect breakpoints and therefore its reliance on predefined target intervals. The model operates at the exon level, analyzing read-depth signals within the boundaries of annotated exonic regions, which constrains its ability to identify novel CNV boundaries or detect structural variants that span multiple exons or extend into intronic regions. This target-based approach, while well-suited for exome sequencing data, limits ExactCN’s capacity to discover unannotated variants or characterize complex structural variations such as inversions, translocations, or intronic duplications that may have functional consequences.

Another important limitation is ExactCN’s inability to perform complete genotyping and determine allelic configurations. The model predicts absolute copy numbers but cannot distinguish between balanced and unbalanced allelic states. For instance, a predicted copy number of 2 could represent either a diploid state (1 copy per allele), a hemizygous duplication (2 copies on one allele, 0 on the other). Similarly, a copy number of 3 could arise from a simple heterozygous duplication (2 + 1) or from an unbalanced state where one allele carries multiple copies. This ambiguity has direct clinical implications and may be critical for some analyses. Resolving such allelic configurations requires haplotype-aware genotyping and phasing information, which is not captured by read-depth signals alone. ExactCN currently does not incorporate additional signals that are commonly used in structural variation detection, such as split reads, discordant read pairs, or B-allele frequencies.

The reliance on DRAGEN calls from matched WGS data as semi-ground truth, while necessary due to the lack of true ground truth, might introduce potential biases into the model’s predictions. Although DRAGEN represents a state-of-the-art CNV calling pipeline with demonstrated high accuracy, any systematic errors or biases in the WGS-based labels may propagate to ExactCN’s learned representations. Future work should aim to validate ExactCN’s predictions using independently verified CNV datasets and orthogonal validation methods. Additionally, future work should aim to extend ExactCN to incorporate additional signals, which we anticipate will enable more complete structural variant characterization, precise breakpoint detection, and haplotype-resolved copy number genotyping, particularly in regions with segmental duplications, low mappability, or complex structural variants.

## Notes

https://zenodo.org/records/17666701

### Competing Interest Statement

The authors have declared no competing interest.

