## Supplementary Material for "ExactCN: Predicting Exact Copy Numbers on Whole Exome Sequencing Data"

### Supplementary Material
### for
### ExactCN: Predicting Exact Copy Numbers on Whole Exome Sequencing Data

#### 1 Supplementary Figures

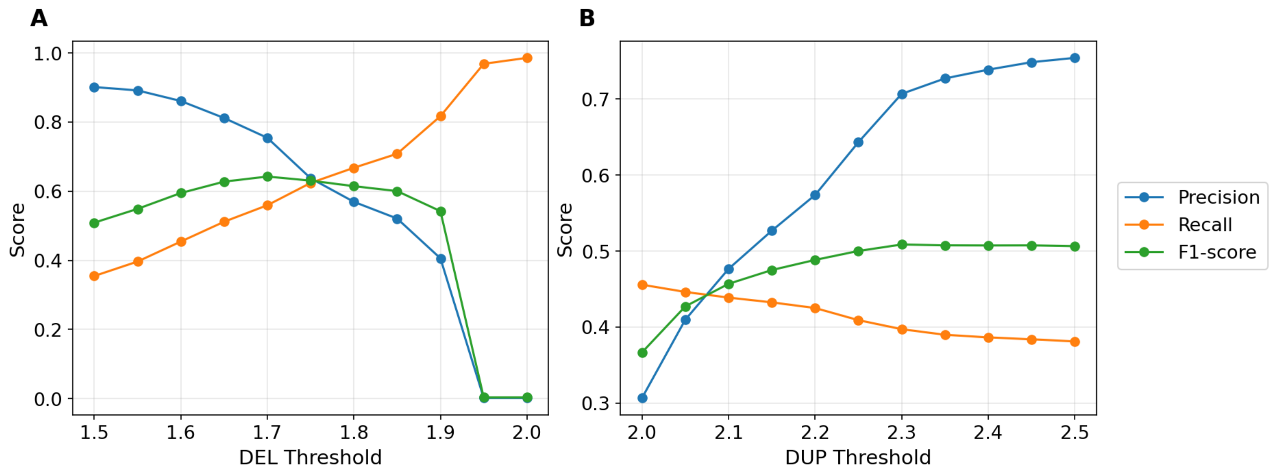


**Supplementary Figure 1.** Performance metrics (Precision, Recall, F1-score) for DEL (A) and DUP (B) detection across threshold values evaluated on 10 validation samples. These thresholds define the classification boundaries: values below the DEL threshold are classified as deletions, values above the DUP threshold are classified as duplications, and values between these thresholds are classified as no call. DEL threshold varied from 1.5 to 2.0 while keeping the DUP threshold fixed at 2.5. DUP threshold varied from 2.0 to 2.5 while keeping the DEL threshold fixed at 1.5.

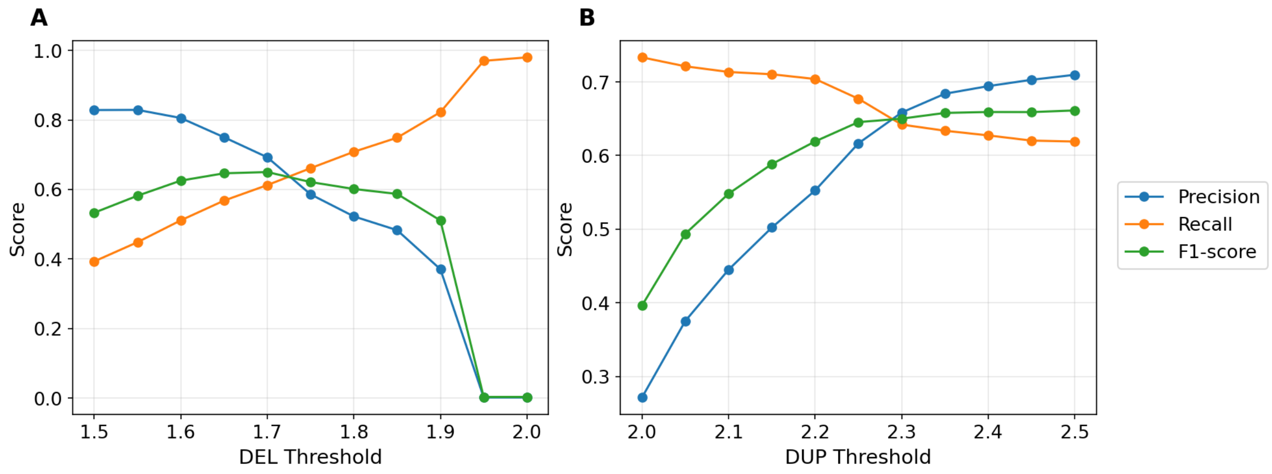


**Supplementary Figure 2.** Performance metrics (Precision, Recall, F1-score) for DEL (A) and DUP (B) detection across threshold values evaluated on 147 test samples. These thresholds define the classification boundaries: values below the DEL threshold are classified as deletions, values above the DUP threshold are classified as duplications, and values between these thresholds are classified as no call. DEL threshold varied from 1.5 to 2.0 while keeping the DUP threshold fixed at 2.5. DUP threshold varied from 2.0 to 2.5 while keeping the DEL threshold fixed at 1.5.

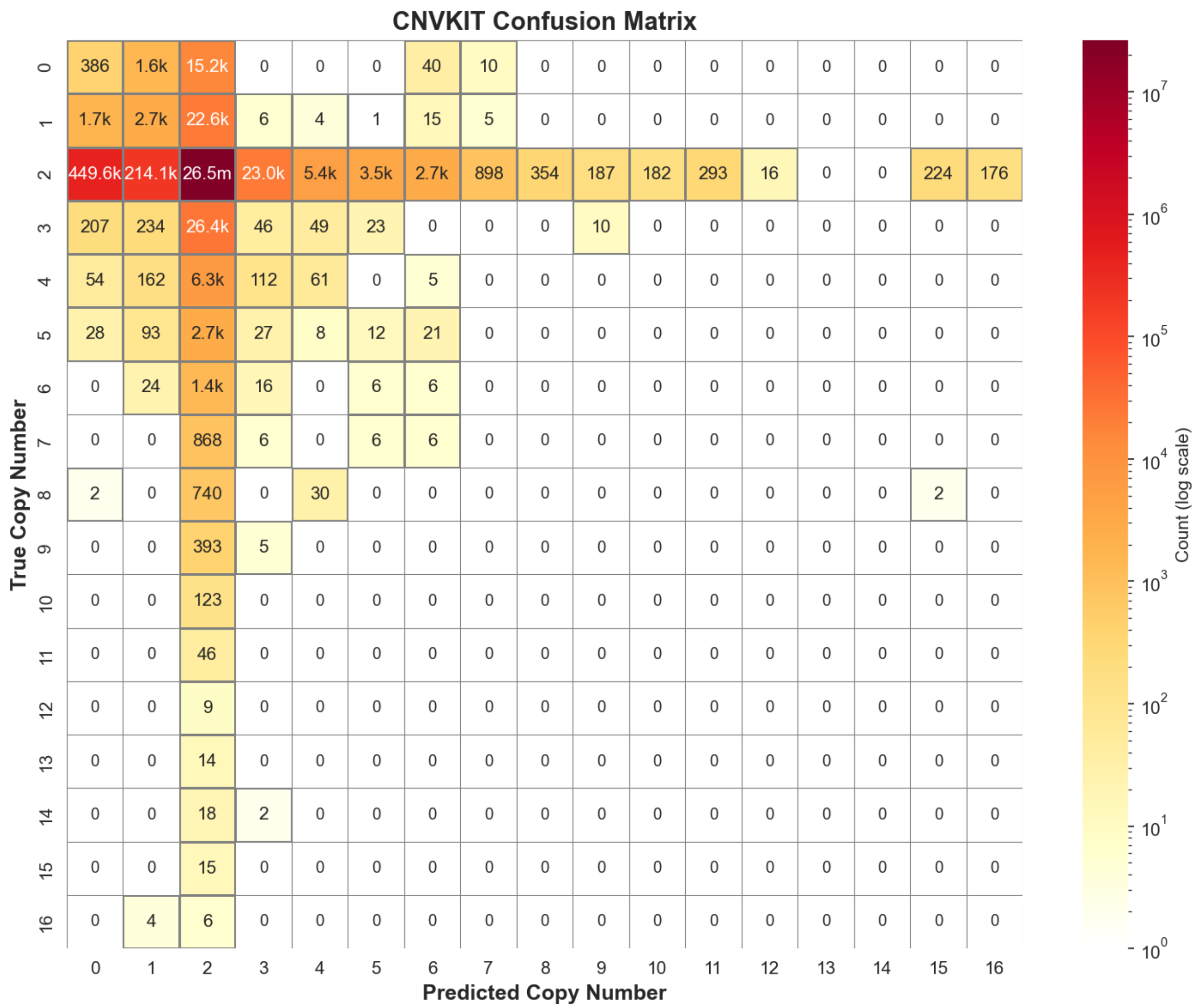


**Supplementary Figure 3.** Confusion matrix for CNVkit copy number predictions. Rows represent true copy numbers and columns represent predicted copy numbers (0-16). Predictions exceeding the maximum observed copy number of 16 are capped at 16. Color intensity indicates the frequency of predictions, with darker colors representing higher counts.

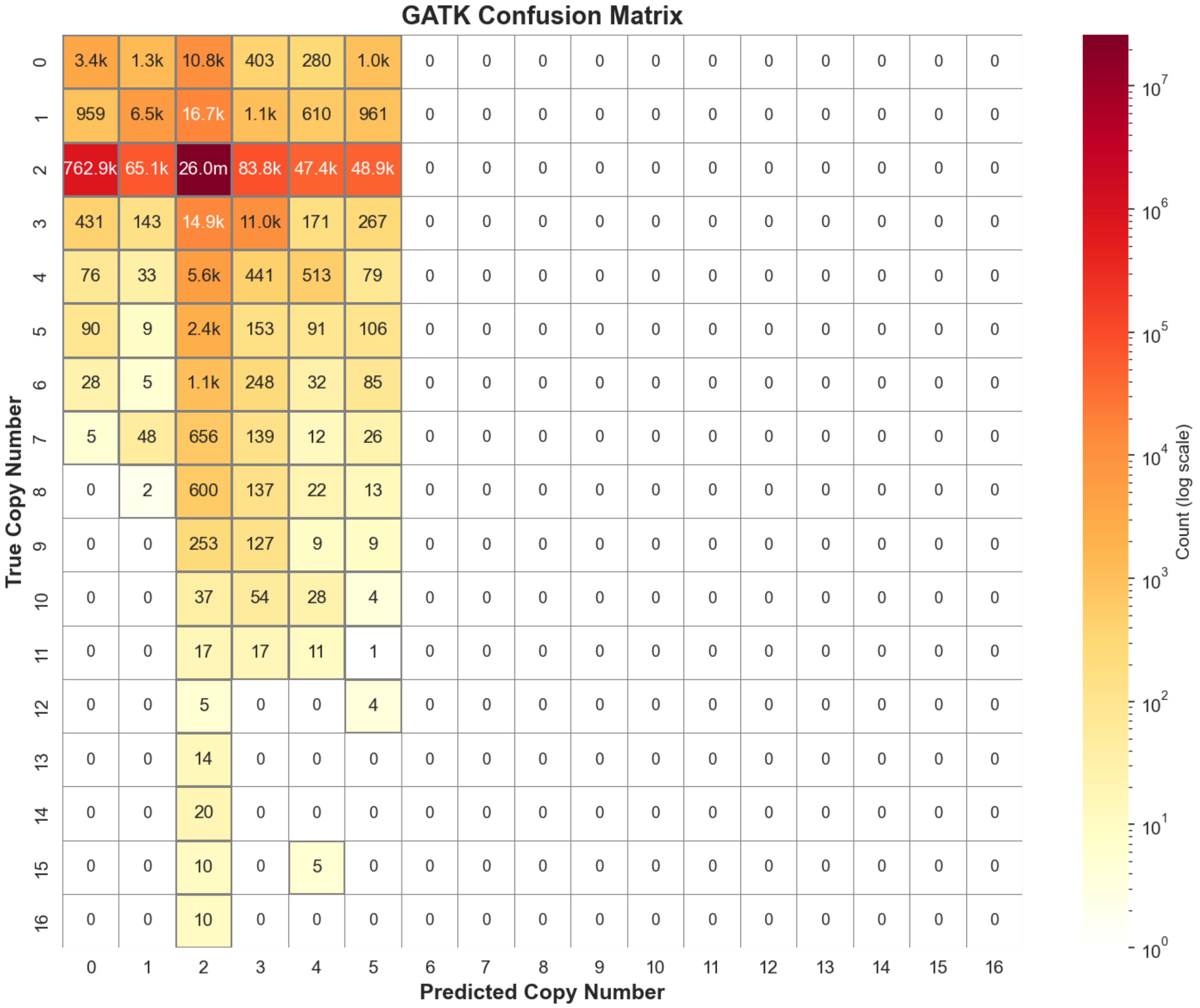


**Supplementary Figure 4.** Confusion matrix for GATK copy number predictions. Rows represent true copy numbers and columns represent predicted copy numbers (0-16). Predictions exceeding the maximum observed copy number of 16 are capped at 16. Color intensity indicates the frequency of predictions, with darker colors representing higher counts.

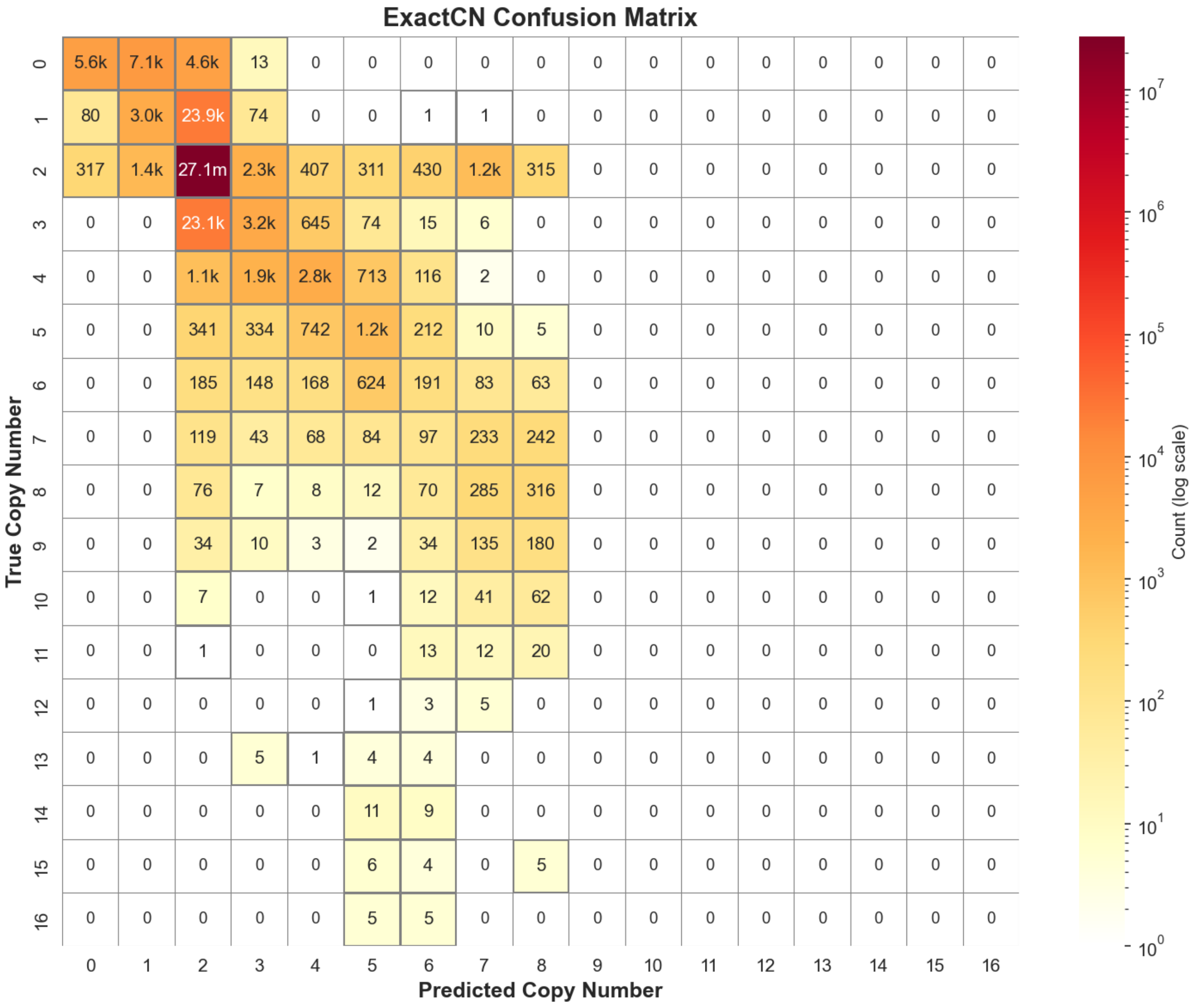


**Supplementary Figure 5.** Confusion matrix for ExactCN copy number predictions. Rows represent true copy numbers and columns represent predicted copy numbers (0-16). Predictions exceeding the maximum observed copy number of 16 are capped at 16. Color intensity indicates the frequency of predictions, with darker colors representing higher counts.

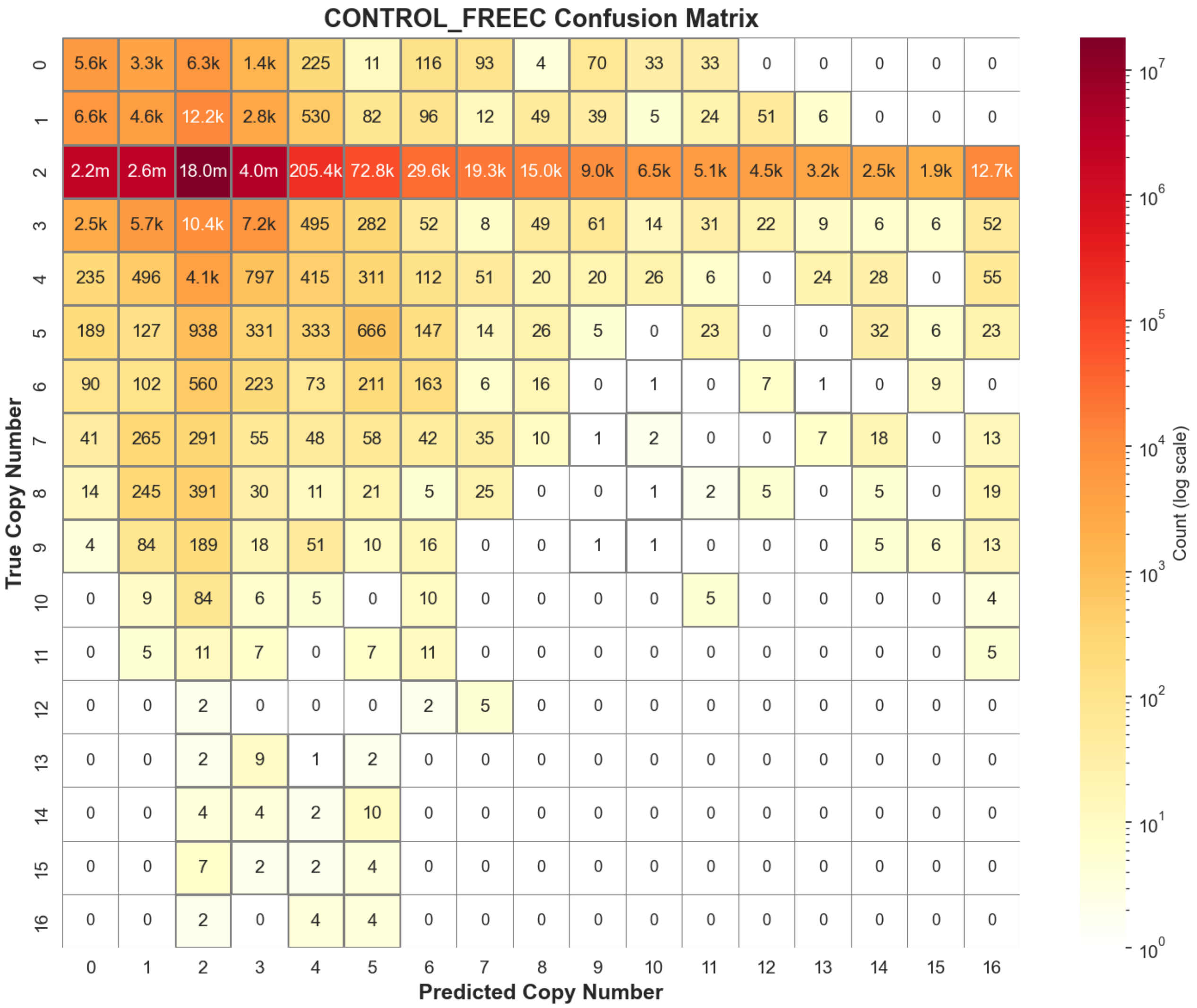


**Supplementary Figure 6.** Confusion matrix for Control-FREEC copy number predictions. Rows represent true copy numbers and columns represent predicted copy numbers (0-16). Predictions exceeding the maximum observed copy number of 16 are capped at 16. Color intensity indicates the frequency of predictions, with darker colors representing higher counts.

#### 2 Supplementary Tables

**Supplementary Table 1.** Extended classification performance metrics including NO-CALL regions. Metrics are calculated for deletions (DEL), duplications (DUP), no-call regions (NO-CALL), and overall performance. Bold values indicate the best performance in each row.

| Metric | CNVkit | Control-FREEC | ECOLE | ECOLE-FT | GATK | ExactCN |
|---|---|---|---|---|---|---|
| **DEL Performance** | | | | | | |
| DEL Precision | 0.009 | 0.004 | 0.230 | **0.816** | 0.014 | 0.755 |
| DEL Recall | 0.145 | 0.454 | 0.419 | **0.636** | 0.277 | 0.559 |
| DEL F1-score | 0.018 | 0.008 | 0.297 | **0.715** | 0.026 | 0.642 |
| **DUP Performance** | | | | | | |
| DUP Precision | 0.010 | 0.003 | 0.104 | 0.551 | 0.067 | **0.713** |
| DUP Recall | 0.012 | 0.319 | 0.371 | **0.425** | 0.334 | 0.413 |
| DUP F1-score | 0.011 | 0.006 | 0.162 | 0.480 | 0.112 | **0.523** |
| **NO-CALL Performance** | | | | | | |
| NO-CALL Precision | 0.997 | 0.998 | 0.998 | **0.999** | 0.998 | 0.998 |
| NO-CALL Recall | 0.974 | 0.662 | 0.993 | 0.999 | 0.961 | **0.999** |
| NO-CALL F1-score | 0.985 | 0.796 | 0.996 | 0.999 | 0.979 | **0.999** |

**Supplementary Table 2.** Additional overall performance metrics. Macro averages treat all classes equally, micro averages weight by support, and weighted averages account for class imbalance. Cohen's Kappa and Quadratic Weighted Kappa (QWK) measure agreement beyond chance. Bold values indicate the best performance in each row.

| Metric | CNVkit | Control-FREEC | ECOLE | ECOLE-FT | GATK | ExactCN |
|---|---|---|---|---|---|---|
| **Macro-Averaged Metrics** | | | | | | |
| Macro Precision | 0.339 | 0.335 | 0.444 | 0.789 | 0.360 | **0.822** |
| Macro Recall | 0.377 | 0.478 | 0.594 | **0.687** | 0.524 | 0.657 |
| Macro F1-score | 0.338 | 0.270 | 0.485 | **0.731** | 0.373 | 0.722 |
| **Micro-Averaged Metrics** | | | | | | |
| Micro Precision | 0.971 | 0.661 | 0.991 | 0.998 | 0.959 | **0.998** |
| Micro Recall | 0.971 | 0.661 | 0.991 | 0.998 | 0.959 | **0.998** |
| Micro F1-score | 0.971 | 0.661 | 0.991 | 0.998 | 0.959 | **0.998** |
| **Weighted-Averaged Metrics** | | | | | | |
| Weighted Precision | 0.994 | 0.995 | 0.996 | **0.998** | 0.995 | 0.998 |
| Weighted Recall | 0.971 | 0.661 | 0.991 | 0.998 | 0.959 | **0.998** |
| Weighted F1-score | 0.982 | 0.794 | 0.993 | **0.998** | 0.976 | 0.998 |
| **Agreement Metrics** | | | | | | |
| Cohen's Kappa | 0.014 | 0.004 | 0.220 | **0.605** | 0.045 | 0.588 |
| Quadratic Weighted Kappa | 0.014 | 0.005 | 0.269 | **0.676** | 0.032 | 0.623 |

**Supplementary Table 3.** Extended regression performance metrics including NO-CALL regions. Mean Absolute Error (MAE) and Root Mean Squared Error (RMSE) are reported for deletions (DEL), duplications (DUP), no-call regions (NO-CALL), and overall performance. Bold values indicate the best performance in each row.

| **Metric** | **CNVkit** | **Control-FREEC** | **GATK** | **ExactCN** |
|---|---|---|---|---|
| **DEL Performance** | | | | |
| DEL MAE | 1.282 | 1.181 | 1.267 | **0.914** |
| DEL RMSE | 1.420 | 1.652 | 1.657 | **1.062** |
| **DUP Performance** | | | | |
| DUP MAE | 1.728 | 1.775 | 1.402 | **0.953** |
| DUP RMSE | 2.215 | 2.556 | 2.077 | **1.239** |
| **NO-CALL Performance** | | | | |
| NO-CALL MAE | 0.044 | 0.458 | 0.072 | **0.001** |
| NO-CALL RMSE | 0.293 | 0.998 | 0.379 | **0.046** |
| **Overall Performance** | | | | |
| Overall MAE | 0.049 | 0.461 | 0.076 | **0.003** |
| Overall RMSE | 0.310 | 1.003 | 0.392 | **0.079** |

**Supplementary Table 4.** Additional regression performance metrics. Macro averages treat all classes equally, weighted averages account for class imbalance. Pearson correlation measures linear relationships, Spearman correlation measures monotonic relationships, and QWK measures ordinal agreement. Bold values indicate the best performance in each row.

| **Metric** | **CNVkit** | **Control-FREEC** | **GATK** | **ExactCN$^{SMN}$** |
|---|---|---|---|---|
| **Macro-Averaged Metrics** | | | | |
| Macro MAE | 1.018 | 1.138 | 0.914 | **0.622** |
| Macro RMSE | 1.309 | 1.735 | 1.371 | **0.783** |
| **Weighted-Averaged Metrics** | | | | |
| Weighted MAE | 0.049 | 0.461 | 0.076 | **0.003** |
| Weighted RMSE | 0.298 | 1.001 | 0.383 | **0.049** |
| **Pearson Correlation** | | | | |
| DEL Pearson | -0.087 | 0.063 | 0.058 | **0.642** |
| DUP Pearson | 0.049 | 0.204 | -0.053 | **0.783** |
| Overall Pearson | 0.014 | 0.028 | 0.032 | **0.669** |
| Macro Pearson | -0.019 | 0.133 | 0.003 | **0.713** |
| **Spearman Correlation** | | | | |
| DEL Spearman | -0.069 | 0.094 | 0.041 | **0.650** |
| DUP Spearman | 0.030 | 0.184 | -0.193 | **0.762** |
| Overall Spearman | 0.024 | 0.021 | 0.068 | **0.550** |
| Macro Spearman | -0.020 | 0.139 | -0.076 | **0.706** |
| **QWK** | | | | |
| Overall | 0.014 | 0.005 | 0.034 | **0.510** |

**Supplementary Table 5.** Classification performance metrics for *SMN1/2* copy number variant detection on 40 WES samples from the 1000 Genomes Project. Chen et al. calls are used as the semi-ground truth to calculate the metrics. Precision, Recall, and F1-score are reported for deletions (DEL), duplications (DUP), no-call regions (NO-CALL), and overall macro-averaged performance. Bold values indicate the best performance in each row.

| **Metric** | **CNVkit** | **Control-FREEC** | **ECOLE** | **ECOLE-FT** | **GATK** | **ExactCN** | **ExactCN$^{SMN}$** |
|---|---|---|---|---|---|---|---|
| **DEL Performance** | | | | | | | |
| DEL Precision | **1.000** | 0.600 | 0.746 | 0.000 | 0.036 | 0.000 | 0.786 |
| DEL Recall | 0.071 | 0.214 | 0.397 | 0.000 | 0.016 | 0.000 | **0.786** |
| DEL F1 | 0.133 | 0.316 | 0.518 | 0.000 | 0.022 | 0.000 | **0.786** |
| **DUP Performance** | | | | | | | |
| DUP Precision | **1.000** | 0.200 | 0.000 | 0.000 | 0.250 | 0.000 | **0.800** |
| DUP Recall | 0.111 | 0.222 | 0.000 | 0.000 | 0.074 | 0.000 | **0.444** |
| DUP F1 | 0.200 | 0.211 | 0.000 | 0.000 | 0.114 | 0.000 | **0.571** |
| **NO-CALL Performance** | | | | | | | |
| NO-CALL Precision | 0.447 | 0.480 | 0.569 | 0.500 | 0.482 | 0.481 | **0.800** |
| NO-CALL Recall | **1.000** | 0.706 | 0.889 | **1.000** | 0.882 | **1.000** | 0.500 |
| NO-CALL F1 | 0.618 | 0.571 | **0.694** | 0.667 | 0.624 | 0.650 | 0.615 |
| **Macro-Averaged Performance** | | | | | | | |
| Macro Precision | 0.816 | 0.427 | 0.438 | 0.167 | 0.256 | 0.160 | **0.795** |
| Macro Recall | 0.394 | 0.381 | 0.429 | 0.333 | 0.324 | 0.333 | **0.577** |
| Macro F1-Score | 0.317 | 0.366 | 0.404 | 0.222 | 0.253 | 0.217 | **0.657** |

**Supplementary Table 6.** Performance metrics for *SMN1/2* copy number regression performances of tools on 40 WES samples from the 1000 Genome Project. Chen et al. calls are used as the semi-ground truth to calculate the metrics. Mean Absolute Error (MAE) and Root Mean Squared Error (RMSE) are reported for deletions (DEL), duplications (DUP), no-call regions (NO-CALL), and overall performance. Bold values indicate the best performance in each row.

| **Metric** | **CNVkit** | **Control-FREEC** | **GATK** | **ExactCN** | **ExactCN$^{SMN}$** |
|---|---|---|---|---|---|
| **DEL Performance** | | | | | |
| DEL MAE | 0.929 | 1.500 | 0.984 | 1.000 | **0.364** |
| DEL RMSE | 0.964 | 2.053 | 0.992 | 1.000 | **0.739** |
| **DUP Performance** | | | | | |
| DUP MAE | 0.889 | 1.778 | 2.383 | 1.000 | **0.200** |
| DUP RMSE | 0.943 | 2.828 | 2.546 | 1.000 | **0.447** |
| **NO-CALL Performance** | | | | | |
| NO-CALL MAE | **0.000** | 0.353 | 0.183 | **0.000** | 0.292 |
| NO-CALL RMSE | **0.000** | 0.686 | 0.560 | **0.000** | 0.540 |
| **Overall Performance** | | | | | |
| Overall MAE | 0.525 | 1.075 | 0.958 | 0.519 | **0.300** |
| Overall RMSE | 0.725 | 1.864 | 1.391 | 0.720 | **0.592** |

**Supplementary Table 7.** Confusion matrices for CNVkit, Control-FREEC, ECOLE, ECOLE-FT, GATK and ExactCN for CNV calling on 147 WES samples.

| | | **Ground Truth** | | |
|---|---|---|---|---|
| TOOLS | **Predicted** | NO-CALL | DUP | DEL |
| CNVKIT | NO-CALL | 26,503,096 | 39,385 | 37,854 |
| | DUP | 40,100 | 469 | 81 |
| | DEL | 667,336 | 810 | 6,417 |
| CONTROL-FREEC | NO-CALL | 18,061,872 | 17,213 | 18,549 |
| | DUP | 4,391,862 | 13,129 | 5,683 |
| | DEL | 4,756,798 | 10,322 | 20,120 |
| ECOLE | NO-CALL | 27,142,839 | 25,891 | 23,821 |
| | DUP | 130,299 | 15,342 | 1,950 |
| | DEL | 61,906 | 163 | 18,581 |
| ECOLE-FT | NO-CALL | 27,190,618 | 23,666 | 15,619 |
| | DUP | 13,665 | 16,901 | 518 |
| | DEL | 6,249 | 97 | 28,215 |
| GATK | NO-CALL | 26,089,636 | 25,926 | 27,486 |
| | DUP | 182,281 | 13,768 | 4,364 |
| | DEL | 841,228 | 887 | 12,217 |
| ExactCN | NO-CALL | 27,135,184 | 23,238 | 23,436 |
| | DUP | 12,822 | 17,127 | 231 |
| | DEL | 3,425 | 7 | 20,395 |

**Supplementary Table 8.** Confusion matrices for CNVkit, Control-FREEC, ECOLE, ECOLE-FT, GATK and ExactCN on SMN1/SMN2 copy number calls.

| | | **Ground Truth** | | |
|---|---|---|---|---|
| TOOLS | **Predicted** | NO-CALL | DUP | DEL |
| CNVKIT | NO-CALL | 17 | 8 | 13 |
| | DUP | 0 | 1 | 0 |
| | DEL | 0 | 0 | 1 |
| CONTROL-FREEC | NO-CALL | 12 | 7 | 6 |
| | DUP | 3 | 2 | 5 |
| | DEL | 2 | 0 | 3 |
| ECOLE | NO-CALL | 17 | 9 | 10 |
| | DUP | 0 | 0 | 0 |
| | DEL | 0 | 0 | 4 |
| ECOLE-FT | NO-CALL | 17 | 9 | 14 |
| | DUP | 0 | 0 | 0 |
| | DEL | 0 | 0 | 0 |
| GATK | NO-CALL | 8 | 0 | 13 |
| | DUP | 9 | 3 | 0 |
| | DEL | 0 | 6 | 1 |
| ExactCN | NO-CALL | 13 | 2 | 12 |
| | DUP | 0 | 0 | 0 |
| | DEL | 0 | 0 | 0 |
| ExactCN$^{SMN}$ | NO-CALL | 14 | 4 | 3 |
| | DUP | 1 | 4 | 0 |
| | DEL | 2 | 1 | 11 |

**Supplementary Table 9.** ECOLE-FT configuration parameters used in this study.

| **Parameter** | **Value** | **Description** |
|---|---|---|
| batch size | 32 | Number of samples processed together before updating weights |
| epochs | 5 | Number of one full pass through the entire training dataset |
| learning rate | 0.00005 | Size of each step taken during weight updates |
| normalization mean | 109.22905835358287 | Average value used to normalize data |
| normalization standard deviation | 101.90851300487935 | The amount data varies around the mean to normalize data |

**Supplementary Table 10.** Control-FREEC configuration parameters used in this study.

| **Parameter** | **Value** | **Description** |
|---|---|---|
| breakPointThreshold | 0.8 | Threshold for breakpoint detection |
| breakPointType | 4 | Type of breakpoint detection algorithm |
| minExpectedGC | 0.35 | Minimum expected GC content |
| maxExpectedGC | 0.55 | Maximum expected GC content |
| degree | 3 | Polynomial degree for GC correction |
| minCNAlength | 1 | Minimum length of copy number alteration (in windows) |
| readCountThreshold | 10 | Minimum read count threshold |
| contaminationAdjustment | FALSE | Adjust for sample contamination |
| forceGCcontentNormalization | 1 | Force GC content normalization |

**Supplementary Table 11.** List of 530 samples from the 1000 Genomes Project selected for pre-training ExactCN.

| | | | | | | | | | |
|---|---|---|---|---|---|---|---|---|---|
| HG00096 | HG00097 | HG00099 | HG00232 | HG00234 | HG00236 | HG00237 | HG00238 | HG00239 | HG00242 |
| HG00245 | HG00246 | HG00250 | HG00251 | HG00252 | HG00253 | HG00254 | HG00255 | HG00256 | HG00257 |
| HG00260 | HG00261 | HG00263 | HG00265 | HG00268 | HG00271 | HG00276 | HG00277 | HG00278 | HG00280 |
| HG00281 | HG00288 | HG00290 | HG00304 | HG00306 | HG00309 | HG00310 | HG00313 | HG00315 | HG00318 |
| HG00321 | HG00324 | HG00325 | HG00326 | HG00327 | HG00328 | HG00329 | HG00330 | HG00331 | HG00332 |
| HG00334 | HG00336 | HG00337 | HG00338 | HG00341 | HG00345 | HG00346 | HG00350 | HG00351 | HG00353 |
| HG00355 | HG00356 | HG00357 | HG00358 | HG00360 | HG00361 | HG00362 | HG00367 | HG00368 | HG00369 |
| HG00372 | HG00373 | HG00375 | HG00376 | HG00378 | HG00379 | HG00382 | HG00384 | HG00403 | HG00404 |
| HG00409 | HG00410 | HG00419 | HG00421 | HG00422 | HG00428 | HG00436 | HG00442 | HG00443 | HG00445 |
| HG00448 | HG00449 | HG00452 | HG00463 | HG00464 | HG00472 | HG00473 | HG00475 | HG00476 | HG00478 |
| HG00479 | HG00500 | HG00525 | HG00531 | HG00534 | HG00536 | HG00537 | HG00542 | HG00543 | HG00551 |
| HG00554 | HG00560 | HG00566 | HG00581 | HG00583 | HG00590 | HG00592 | HG00593 | HG00595 | HG00596 |
| HG00598 | HG00599 | HG00607 | HG00610 | HG00611 | HG00614 | HG00619 | HG00622 | HG00623 | HG00625 |
| HG00628 | HG00629 | HG00631 | HG00632 | HG00634 | HG00640 | HG00653 | HG00656 | HG00657 | HG00662 |
| HG00663 | HG00672 | HG00674 | HG00684 | HG00689 | HG00690 | HG00692 | HG00698 | HG00699 | HG00701 |
| HG00732 | HG00734 | HG00736 | HG00740 | HG00743 | HG00766 | HG00844 | HG00864 | HG00867 | HG00879 |
| HG00956 | HG00978 | HG01029 | HG01031 | HG01046 | HG01047 | HG01048 | HG01049 | HG01051 | HG01055 |
| HG01058 | HG01061 | HG01063 | HG01066 | HG01069 | HG01072 | HG01073 | HG01075 | HG01077 | HG01079 |
| HG01085 | HG01086 | HG01088 | HG01092 | HG01094 | HG01095 | HG01101 | HG01104 | HG01105 | HG01107 |
| HG01108 | HG01110 | HG01111 | HG01112 | HG01113 | HG01119 | HG01121 | HG01124 | HG01125 | HG01130 |
| HG01131 | HG01133 | HG01134 | HG01136 | HG01137 | HG01139 | HG01140 | HG01142 | HG01148 | HG01149 |
| HG01161 | HG01162 | HG01164 | HG01168 | HG01171 | HG01173 | HG01174 | HG01176 | HG01182 | HG01187 |
| HG01190 | HG01191 | HG01197 | HG01198 | HG01200 | HG01204 | HG01205 | HG01241 | HG01247 | HG01248 |
| HG01250 | HG01253 | HG01256 | HG01257 | HG01259 | HG01269 | HG01271 | HG01272 | HG01281 | HG01284 |
| HG01302 | HG01305 | HG01308 | HG01323 | HG01326 | HG01334 | HG01341 | HG01342 | HG01344 | HG01345 |
| HG01348 | HG01350 | HG01351 | HG01353 | HG01354 | HG01356 | HG01357 | HG01359 | HG01360 | HG01362 |
| HG01363 | HG01365 | HG01366 | HG01369 | HG01372 | HG01375 | HG01384 | HG01389 | HG01393 | HG01396 |
| HG01398 | HG01402 | HG01412 | HG01413 | HG01432 | HG01437 | HG01438 | HG01440 | HG01441 | HG01444 |
| HG01447 | HG01455 | HG01456 | HG01459 | HG01461 | HG01462 | HG01468 | HG01474 | HG01485 | HG01488 |
| HG01489 | HG01491 | HG01492 | HG01494 | HG01495 | HG01497 | HG01501 | HG01503 | HG01504 | HG01506 |
| HG01507 | HG01509 | HG01512 | HG01516 | HG01518 | HG01521 | HG01524 | HG01525 | HG01530 | HG01531 |
| HG01537 | HG01550 | HG01551 | HG01556 | HG01565 | HG01566 | HG01571 | HG01572 | HG01578 | HG01583 |
| HG01589 | HG01593 | HG01595 | HG01597 | HG01598 | HG01600 | HG01603 | HG01605 | HG01606 | HG01610 |
| HG01612 | HG01613 | HG01617 | HG01620 | HG01623 | HG01625 | HG01628 | HG01630 | HG01631 | HG01668 |
| HG01670 | HG01673 | HG01682 | HG01684 | HG01686 | HG01694 | HG01699 | HG01700 | HG01702 | HG01704 |
| HG01708 | HG01709 | HG01710 | HG01747 | HG01756 | HG01757 | HG01762 | HG01765 | HG01768 | HG01771 |
| HG01776 | HG01777 | HG01779 | HG01783 | HG01789 | HG01790 | HG01794 | HG01797 | HG01798 | HG01800 |
| HG01801 | HG01802 | HG01805 | HG01806 | HG01809 | HG01810 | HG01811 | HG01812 | HG01815 | HG01816 |
| HG01817 | HG01840 | HG01844 | HG01845 | HG01847 | HG01848 | HG01849 | HG01851 | HG01855 | HG01857 |
| HG01858 | HG01859 | HG01860 | HG01863 | HG01865 | HG01866 | HG01868 | HG01869 | HG01870 | HG01871 |
| HG01879 | HG01880 | HG01882 | HG01883 | HG01889 | HG01890 | HG01893 | HG01894 | HG01912 | HG01914 |
| HG01915 | HG01918 | HG01920 | HG01921 | HG01924 | HG01926 | HG01927 | HG01932 | HG01935 | HG01936 |
| HG01938 | HG01941 | HG01945 | HG01950 | HG01951 | HG01953 | HG01956 | HG01958 | HG01965 | HG01968 |
| HG01970 | HG01971 | HG01974 | HG01976 | HG01979 | HG01980 | HG01988 | HG01990 | HG01991 | HG01992 |
| HG02002 | HG02006 | HG02009 | HG02016 | HG02017 | HG02019 | HG02023 | HG02028 | HG02031 | HG02040 |
| HG02047 | HG02048 | HG02050 | HG02051 | HG02053 | HG02054 | HG02057 | HG02058 | HG02060 | HG02061 |
| HG02064 | HG02069 | HG02070 | HG02072 | HG02073 | HG02075 | HG02076 | HG02079 | HG02081 | HG02082 |
| HG02084 | HG02085 | HG02086 | HG02088 | HG02089 | HG02090 | HG02095 | HG02102 | HG02104 | HG02113 |
| HG02116 | HG02128 | HG02134 | HG02136 | HG02137 | HG02138 | HG02139 | HG02140 | HG02142 | HG02143 |
| HG02147 | HG02151 | HG02152 | HG02153 | HG02155 | HG02156 | HG02164 | HG02165 | HG02166 | HG02179 |
| HG02182 | HG02185 | HG02186 | HG02187 | HG02188 | HG02190 | HG02215 | HG02219 | HG02220 | HG02223 |
| HG02230 | HG02231 | HG02232 | HG02233 | HG02238 | HG02239 | HG02250 | HG02256 | HG02259 | HG02262 |
| HG02265 | HG02266 | HG02272 | HG02277 | HG02278 | HG02282 | HG02283 | HG02284 | HG02285 | HG02286 |

**Supplementary Table 12.** List of 10 samples from the 1000 Genomes Project selected for the validation of ExactCN.

| | | | | | | | | | |
|---|---|---|---|---|---|---|---|---|---|
| HG00233 | HG00235 | HG00240 | HG00258 | HG00262 | HG00264 | HG00267 | HG00269 | HG00273 | HG00284 |

**Supplementary Table 13.** List of 147 samples from the 1000 Genomes Project selected for testing ExactCN.

| | | | | | | | | | |
|---|---|---|---|---|---|---|---|---|---|
| HG00285 | HG00311 | HG00319 | HG00335 | HG00343 | HG00366 | HG00371 | HG00380 | HG00406 | HG00451 |
| HG00457 | HG00458 | HG00513 | HG00524 | HG00553 | HG00556 | HG00557 | HG00559 | HG00565 | HG00580 |
| HG00584 | HG00589 | HG00608 | HG00620 | HG00626 | HG00638 | HG00650 | HG00651 | HG00671 | HG00683 |
| HG00705 | HG00708 | HG00728 | HG00729 | HG00739 | HG00881 | HG00982 | HG01028 | HG01054 | HG01060 |
| HG01067 | HG01083 | HG01097 | HG01102 | HG01122 | HG01167 | HG01183 | HG01188 | HG01242 | HG01254 |
| HG01260 | HG01277 | HG01311 | HG01312 | HG01325 | HG01374 | HG01377 | HG01395 | HG01403 | HG01405 |
| HG01414 | HG01435 | HG01443 | HG01465 | HG01479 | HG01486 | HG01498 | HG01510 | HG01522 | HG01527 |
| HG01528 | HG01586 | HG01596 | HG01599 | HG01602 | HG01618 | HG01624 | HG01669 | HG01678 | HG01679 |
| HG01685 | HG01697 | HG01705 | HG01707 | HG01766 | HG01767 | HG01770 | HG01775 | HG01781 | HG01784 |
| HG01799 | HG01807 | HG01813 | HG01841 | HG01842 | HG01843 | HG01852 | HG01862 | HG01864 | HG01874 |
| HG01878 | HG01885 | HG01892 | HG01896 | HG01917 | HG01923 | HG01933 | HG01947 | HG01948 | HG01954 |
| HG01961 | HG01967 | HG01985 | HG01986 | HG01997 | HG02008 | HG02010 | HG02013 | HG02014 | HG02020 |
| HG02025 | HG02029 | HG02035 | HG02052 | HG02067 | HG02078 | HG02087 | HG02105 | HG02107 | HG02108 |
| HG02130 | HG02133 | HG02144 | HG02146 | HG02180 | HG02181 | HG02184 | HG02221 | HG02252 | HG02255 |
| HG02260 | HG02274 | HG02281 | HG02304 | HG02318 | HG02323 | HG02348 | | | |

**Supplementary Table 14.** List of 65 samples from the 1000 Genomes Project selected for ECOLE fine-tuning

| | | | | | | | | | |
|---|---|---|---|---|---|---|---|---|---|
| HG00232 | HG00234 | HG00245 | HG00331 | HG00338 | HG00355 | HG00367 | HG00369 | HG00379 | HG00403 |
| HG00409 | HG00443 | HG00476 | HG00534 | HG00551 | HG00554 | HG00672 | HG00734 | HG01049 | HG01063 |
| HG01066 | HG01077 | HG01079 | HG01086 | HG01124 | HG01131 | HG01140 | HG01182 | HG01247 | HG01257 |
| HG01326 | HG01362 | HG01369 | HG01389 | HG01432 | HG01462 | HG01489 | HG01503 | HG01551 | HG01595 |
| HG01605 | HG01625 | HG01628 | HG01694 | HG01700 | HG01709 | HG01771 | HG01809 | HG01810 | HG01879 |
| HG01938 | HG02053 | HG02057 | HG02060 | HG02072 | HG02116 | HG02139 | HG02156 | HG02164 | HG02165 |
| HG02223 | HG02232 | HG02256 | HG02272 | HG02322 | | | | | |

**Supplementary Table 15.** List of samples from the 1000 Genomes dataset used to fine-tune and test the performance on the SMN1 and SMN2 genes.

| Fine-tune Samples | | | Test Samples | | |
|---|---|---|---|---|---|
| **Sample** | **SMN1 Label** | **SMN2 Label** | **Sample** | **SMN1 Label** | **SMN2 Label** |
| HG00096 | NO-CALL | DEL | HG00233 | NO-CALL | DUP |
| HG00097 | NO-CALL | NO-CALL | HG00235 | NO-CALL | NO-CALL |
| HG00099 | NO-CALL | DEL | HG00240 | NO-CALL | NO-CALL |
| HG00232 | NO-CALL | NO-CALL | HG00262 | NO-CALL | DEL |
| HG00234 | NO-CALL | NO-CALL | HG00267 | NO-CALL | DEL |
| HG00236 | NO-CALL | DEL | HG00273 | NO-CALL | NO-CALL |
| HG00245 | NO-CALL | NO-CALL | HG00284 | NO-CALL | NO-CALL |
| HG00246 | NO-CALL | NO-CALL | HG00285 | NO-CALL | NO-CALL |
| HG00251 | NO-CALL | NO-CALL | HG00311 | NO-CALL | DEL |
| HG00252 | NO-CALL | DEL | HG00329 | NO-CALL | DUP |
| HG00253 | NO-CALL | NO-CALL | HG00335 | NO-CALL | NO-CALL |
| HG00254 | NO-CALL | DEL | HG00380 | NO-CALL | DEL |
| HG00256 | NO-CALL | DEL | HG00406 | NO-CALL | NO-CALL |
| HG00257 | NO-CALL | DEL | HG00451 | NO-CALL | NO-CALL |
| HG00260 | NO-CALL | DEL | HG00457 | NO-CALL | DEL |
| HG00261 | NO-CALL | NO-CALL | HG00458 | NO-CALL | DEL |
| HG00263 | NO-CALL | NO-CALL | HG00524 | NO-CALL | NO-CALL |
| HG00265 | NO-CALL | DEL | HG00553 | NO-CALL | DEL |
| HG00271 | NO-CALL | DEL | HG00565 | NO-CALL | DEL |
| HG00276 | NO-CALL | NO-CALL | HG00566 | DUP | NO-CALL |
| HG00277 | NO-CALL | DEL | HG00580 | NO-CALL | NO-CALL |
| HG00278 | NO-CALL | NO-CALL | HG00584 | NO-CALL | DEL |
| HG00280 | NO-CALL | DEL | HG00589 | NO-CALL | NO-CALL |
| HG00288 | NO-CALL | NO-CALL | HG00626 | NO-CALL | DUP |
| HG00315 | NO-CALL | NO-CALL | HG00651 | NO-CALL | NO-CALL |
| HG00318 | NO-CALL | DUP | HG00708 | NO-CALL | NO-CALL |
| HG00324 | DEL | NO-CALL | HG00732 | NO-CALL | DUP |
| HG00325 | NO-CALL | NO-CALL | HG00739 | NO-CALL | NO-CALL |
| HG00327 | NO-CALL | NO-CALL | HG01061 | NO-CALL | DUP |
| HG00330 | NO-CALL | NO-CALL | HG01183 | NO-CALL | NO-CALL |
| HG00331 | NO-CALL | NO-CALL | HG01707 | NO-CALL | NO-CALL |
| HG01085 | DEL | NO-CALL | HG01892 | DEL | DEL |
| HG01094 | NO-CALL | DUP | HG01896 | NO-CALL | DEL |
| HG01205 | DEL | DEL | HG01948 | DEL | NO-CALL |
| HG01241 | DUP | NO-CALL | HG02013 | NO-CALL | NO-CALL |
| HG01393 | NO-CALL | DUP | HG02089 | DUP | NO-CALL |
| HG01447 | DUP | NO-CALL | HG02128 | NO-CALL | DUP |
| HG01455 | DEL | NO-CALL | HG02180 | DEL | DEL |
| HG01612 | DEL | NO-CALL | HG02255 | DUP | DUP |
| HG01623 | NO-CALL | NO-CALL | HG02281 | NO-CALL | DEL |
| HG01628 | NO-CALL | NO-CALL | | | |
| HG01684 | DUP | NO-CALL | | | |
| HG01686 | DEL | DEL | | | |
| HG01747 | NO-CALL | DUP | | | |
| HG01765 | NO-CALL | NO-CALL | | | |
| HG01801 | DEL | DEL | | | |
| HG01860 | NO-CALL | DUP | | | |
| HG01863 | DEL | NO-CALL | | | |
| HG01914 | NO-CALL | DUP | | | |
| HG01965 | DUP | NO-CALL | | | |
| HG01971 | NO-CALL | DUP | | | |
| HG02051 | NO-CALL | DUP | | | |
| HG02070 | NO-CALL | DUP | | | |
| HG02187 | NO-CALL | NO-CALL | | | |
| HG02262 | NO-CALL | DUP | | | |
